# Pathologic gene network rewiring implicates PPP1R3A as a central cardioprotective factor in pressure overload heart failure

**DOI:** 10.1101/038174

**Authors:** Pablo Cordero, Victoria N. Parikh, Ayca Erbilgin, Ching Shang, Kevin S. Smith, Frederick Dewey, Kathia Zaleta, Michael Morley, Jeff Brandimarto, Nicole Glazer, Aleksandra Pavlovic, Christine Moravec, Wilson H. Tang, Jamie Viterna, Christine Malloy, Sridhar Hannenhalli, Hongzhe Li, Scott Ritter, Mingyao Li, Andrew Connolly, Hakon Hakonarson, Aldons J. Lusis, Kenneth B. Margulies, Anna A. Depaoli-Roach, Stephen Montgomery, Matthew T. Wheeler, Thomas Cappola, Euan A. Ashley

## Abstract

Heart failure is a leading cause of mortality, yet our understanding of the genetic interactions underlying this disease remains incomplete. Here, we harvested 1352 healthy and failing human hearts directly from transplant center operating rooms, and obtained genome-wide genotyping and gene expression measurements for a subset of 313. We built failing and non-failing cardiac regulatory gene networks, revealing important regulators and cardiac expression quantitative trait loci (eQTLs). *PPP1R3A* emerged as a novel regulator whose network connectivity changed significantly between health and disease. Time-course RNA sequencing after *PPP1R3A* knock-down validated network-based predictions of metabolic pathway expression, increased cardiomyocyte size, and perturbed respiratory metabolism. Mice lacking *PPP1R3A* were protected against pressure-overload heart failure. We present a global gene interaction map of the human heart failure transition, identify new cardiac eQTLs, and demonstrate the discovery potential of disease-specific networks through the description of *PPP1R3A* as a novel central protective regulator in heart failure.

---

Heart failure (HF) is a life-threatening syndrome characterized by an inability of the heart to meet the metabolic demands of the body and costs the US more than $34 billion a year to treat 6 million patients.^1,2^ Despite this, the underlying molecular mechanisms remain poorly understood and the few approved therapeutics target maladaptive compensatory mechanisms rather than proximate molecular mechanisms ^3,4^.

Studies of gene expression in heart failure have focused predominantly on determining transcriptional signatures in small numbers of diseased human hearts or have used animal models to examine changes in established pathways^5–9^. These efforts have revealed changes in gene expression of key sarcomeric, calcium cycling, and metabolic genes. However, due to the lack of a comprehensive gene regulatory network of the failing heart, it remains unclear how these differentially expressed genes interact and to what extent they are causal for disease. The lack of expansive cardiac gene expression measurements in sufficient numbers of human tissues has precluded the creation of an interaction map of the failing heart. In addition, because of the significant logistical challenge of harvesting healthy hearts, very few studies have included a non-failing control group, making conclusions regarding the transition to heart failure hard to draw. Finally, the high metabolic rate of the heart limits the utility of tissue collected post-mortem, such as that from public resources such as GTEx ^10–12^, since gene expression programs are rapidly altered in an environment of high oxidative and nitrosative stress ^13^.

## Results

### Real time harvesting of transplant hearts yields high-quality transcriptomic measurements

The MAGnet consortium was founded to establish best practices for the harvesting of human cardiac tissue (see Online Methods) and to explore the genetic landscape of cardiac gene expression ^7,13,14^. Using this consensus protocol, we obtained 1352 human cardiac samples and chose a subset of 313 hearts, including 177 failing hearts collected immediately post-transplantation and 136 healthy donor controls that were suitable for transplantation but did not reach a recipient due to logistical reasons. We measured left-ventricular genome-wide gene expression and genotyping of these samples and corrected the measurements for known covariates (see Supplementary Methods). We assessed the quality of these measurements in several ways. First, we found that disease status was the dominant source of variation suggesting no major confounding sources of variation (Figure 1A). Second, we confirmed enhanced expression of *NPPA* and *NPPB*, depletion of *SERCA2A*, and isoform switch from *MYH7* towards *MYH6* expression -- established signatures of heart failure (Supplemental Figures 1A and B). Finally, since our sample collection was done on-site immediately before or after cardiac transplantation, unlike post-mortem samples, we were able to investigate whether gene programs involved with oxidative stress were unperturbed. We compared oxidative stress gene expression in our samples to left ventricular sample data in GTEX, collected post-mortem, (data obtained from the recount2 database ^15^) and found that our samples conserved comparable contractility gene expression but had significantly less oxidative gene expression and less perturbation in other metabolic pathways (Supplemental Figure 1C). Having established the quality of our data, we limited our downstream analyses to the top 40% most variable genes (n=7960).

**Figure 1.**
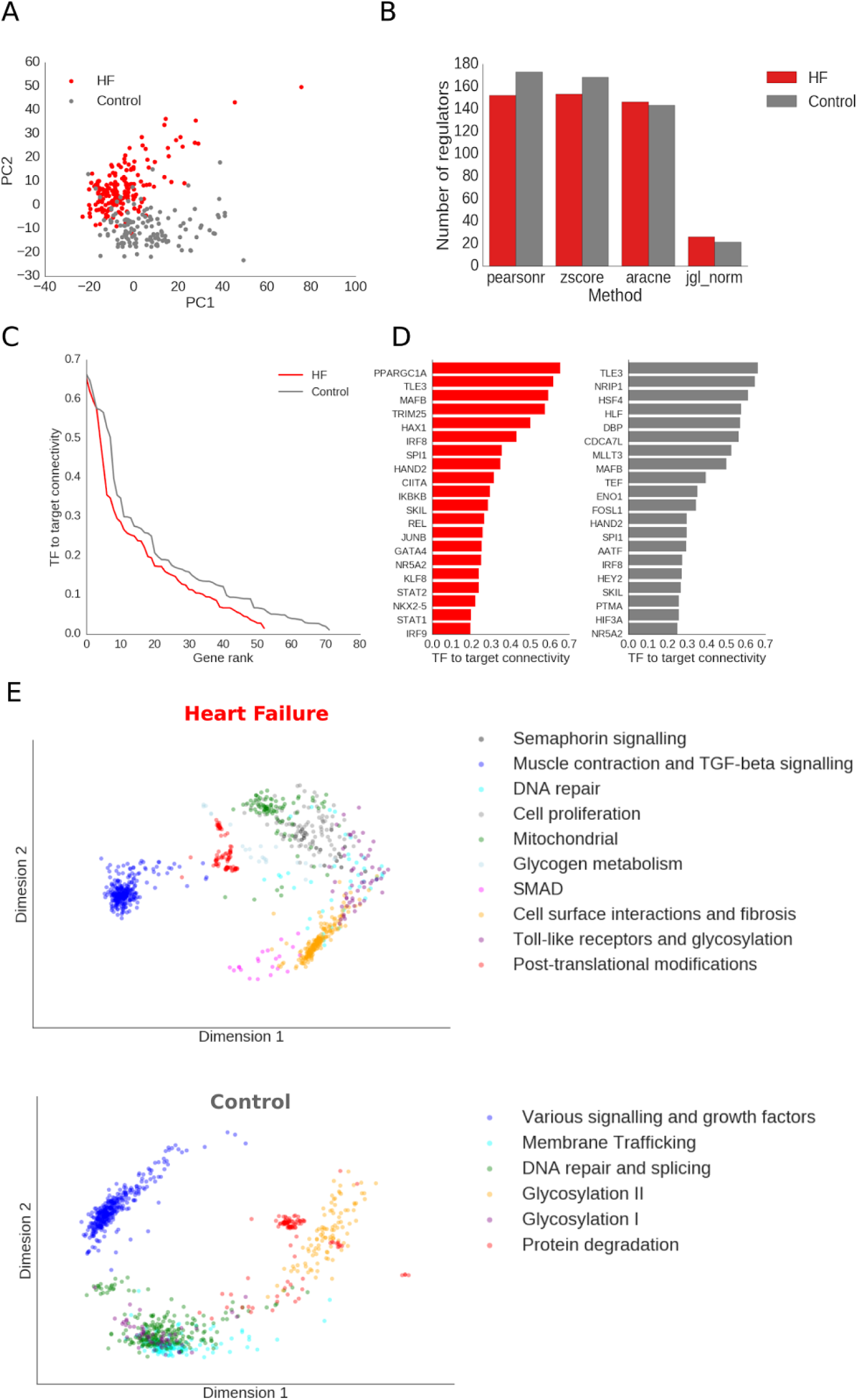
Regulatory rewiring of co-expression networks in heart failure. (A) Principal component analysis of gene expression profiles for 177 failing hearts and 136 non-failing, control, hearts. (B) Number of transcription factors that were deemed significantly more connected than average for several methods for network inference: ARACNE, Z-score, Pearson correlation, and the joint graphical LASSO. Overall, more transcription factors have higher connectivity in the control hearts and Pearson correlation detects the most. (C) Transcription factor to target connectivity scores of known human transcription factors in the failing and control networks. (D). Most active transcription factors as assessed by their co-expression with their respective targets for failing (red) and control (gray) networks. (E) Two-dimensional projection of highly connected subnetworks of genes in the failing and control networks using node2vec, clustered into gene modules.

### The cardiac gene co-expression map exhibits a drastic change of regulators in heart failure

We inferred undirected, cohort-specific gene co-expression networks. Gene regulatory network inference from co-expression is a challenging problem that no single method solves adequately in all contexts. Therefore, we tested four widely used methods: ARACNE ^16^, inverse covariance estimation via the Joint Graphical LASSO (JGL) ^17^, the ZScore method ^18^, and the standard gene-by-gene Pearson correlation. To evaluate the resulting networks, we looked at the connectivity patterns of known transcription factors. Given their role in transcriptional regulation, we expected them to be more connected to other genes than most genes. We therefore tested how many known transcription factors were significantly connected in all networks and all methods (Z test, FDR cutoff 0.05 using the Benjamini-Hochberg method). While all methods showed significant connectivity for similar numbers of transcription factors, Pearson correlation yielded the combined highest number of transcription factor regulators (Figure 1B). We therefore chose the Pearson correlation networks for subsequent analyses.

Between the heart failure and non-failing control networks, known regulators changed significantly. The number of regulators that were highly connected was greater in the control network (Figure 1D) and the top 20 most connected regulators were different between networks (Figure 1E). We found that the highly connected regulators of both networks had known co-expression, physical interaction, and shared protein domain relationships (Supplemental Figure 2B).

Both networks formed cohesive gene modules as apparent through a node embedding visualization ^19^ and Birch clustering (Figure 1E). Each group of genes was specifically enriched with functional annotations as revealed by enrichR ^20^, with the heart failure modules having more diversity of signaling and metabolic annotations.

We then manually curated sets of genes representing five key processes and cellular structures involved in heart failure (metabolic, sarcomeric, electric-contraction (EC) coupling, cell adhesion, and cardiac remodeling). Network connectivity changed between both networks: the failing heart network saw a general rewiring in connectivity within and without these gene sets (Figure 2), with EC coupling genes gaining connectivity with cell adhesion genes but losing connectivity with sarcomeric, and cardiac remodeling (Figure 2A, red); metabolic genes gaining a few specific ‘hubs’ such as the protein phosphatase 1 catalytic and regulatory subunits (*PPP1CC, PPP1R1A*, and *PPP1R3A/B/C*) and the muscle 6-phosphofructokinase *PFKM* in the heart failure network (Figure 2B, blue); cell adhesion gene observing increase in connectivity within its own genes but loss of connectivity with others in heart failure (Figure 2C, purple); and sarcomeric and cardiac remodeling genes observing an increase in connectivity throughout other processes (Figure 2D and 2E orange and green, respectively).

**Figure 2.**
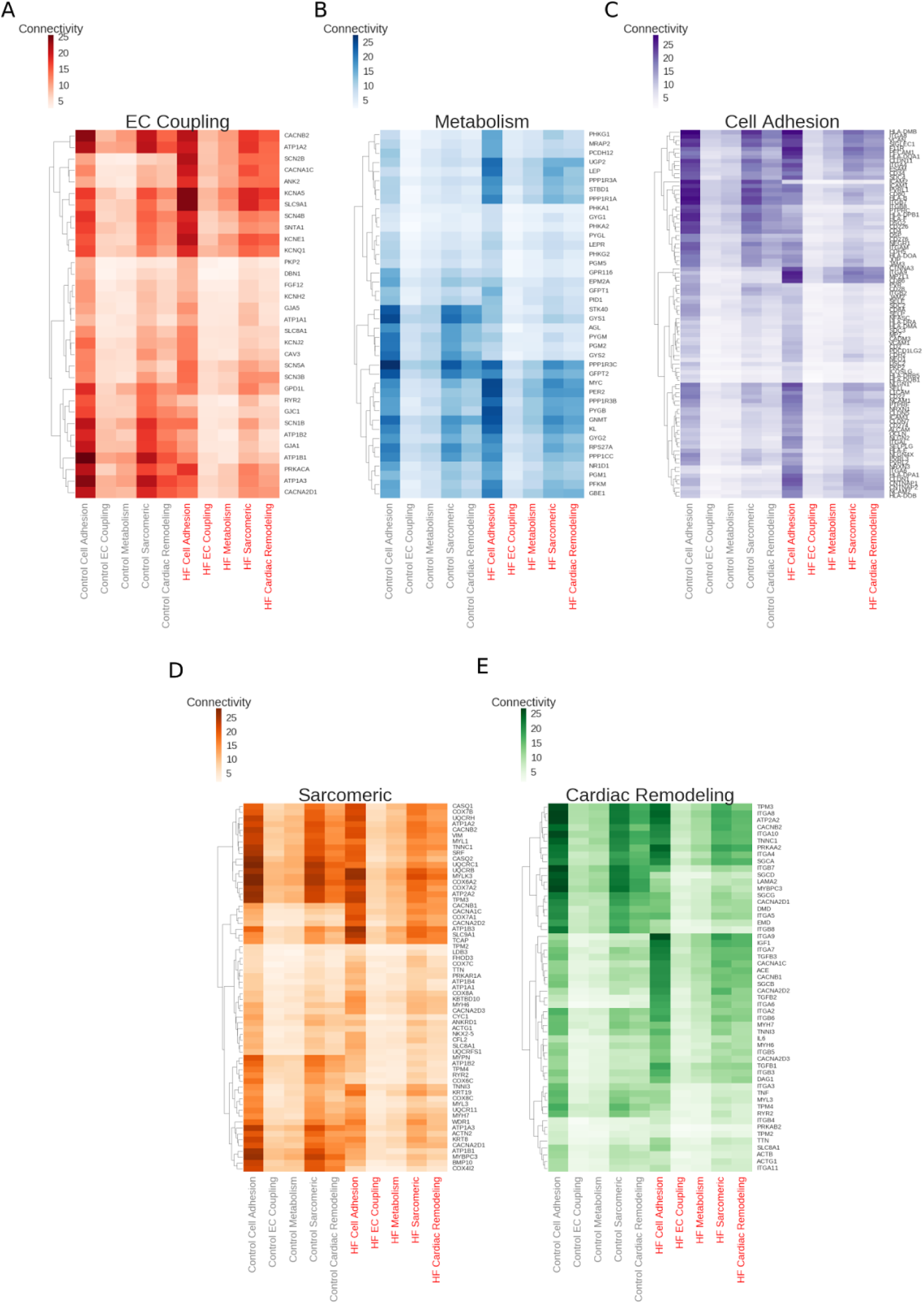
Differential connectivity of known biological processes in heart failure. (A) Mean correlation between five known processes that play critical roles in heart failure: cardiac remodeling, metabolism, cell adhesion, sarcomeric genes, and contractility in the failing and control networks. (B-F) Change in gene-by-gene degree breakdown for cardiac remodeling, sarcomeric, cell adhesion, contractility, and metabolism genes.

### Network-enhanced expression profiles reveal novel cis and trans cardiac eQTLs

We then leveraged genome-wide genotypes to find gene-expression-controlling loci (eQTLs) in each cohort. To increase power for finding significant eQTLs, we applied graph convolution to the gene expression measurements prior to association testing (Supplementary Methods). We then used QTLtools to perform association testing for each cohort separately (Online Methods). We found that the heart failure cohort had more associated eQTLs than the control group (867 vs 416, respectively, Supplementary Files S1 and S2); as expected these eQTLs showed proximity to known transcription factor binding sites (Figure 3A). We then tested these eQTLs for enrichment of regulatory associations using RegulomeDB, a database of known and predicted regulatory regions of the genome ^21^. Here again, the heart failure cohort had higher number of eQTLs with regulatory annotation than the control group (356/867 [43%] and 165/416 [40%] of variants, for failing vs control, respectively, had some adjacent regulatory signature and were predicted for transcription factor binding (Figure S3). We then compared our eQTLs with those found by the GTEx project. Our set of eQTLs contained hundreds of novel assoaciations while overlapping known associations for cardiac left ventricular tissue (431/867 [50% novel associations] for the heart failure group and 141/416 [34% novel associations] for the control group;Figure 3B).

**Figure 3.**
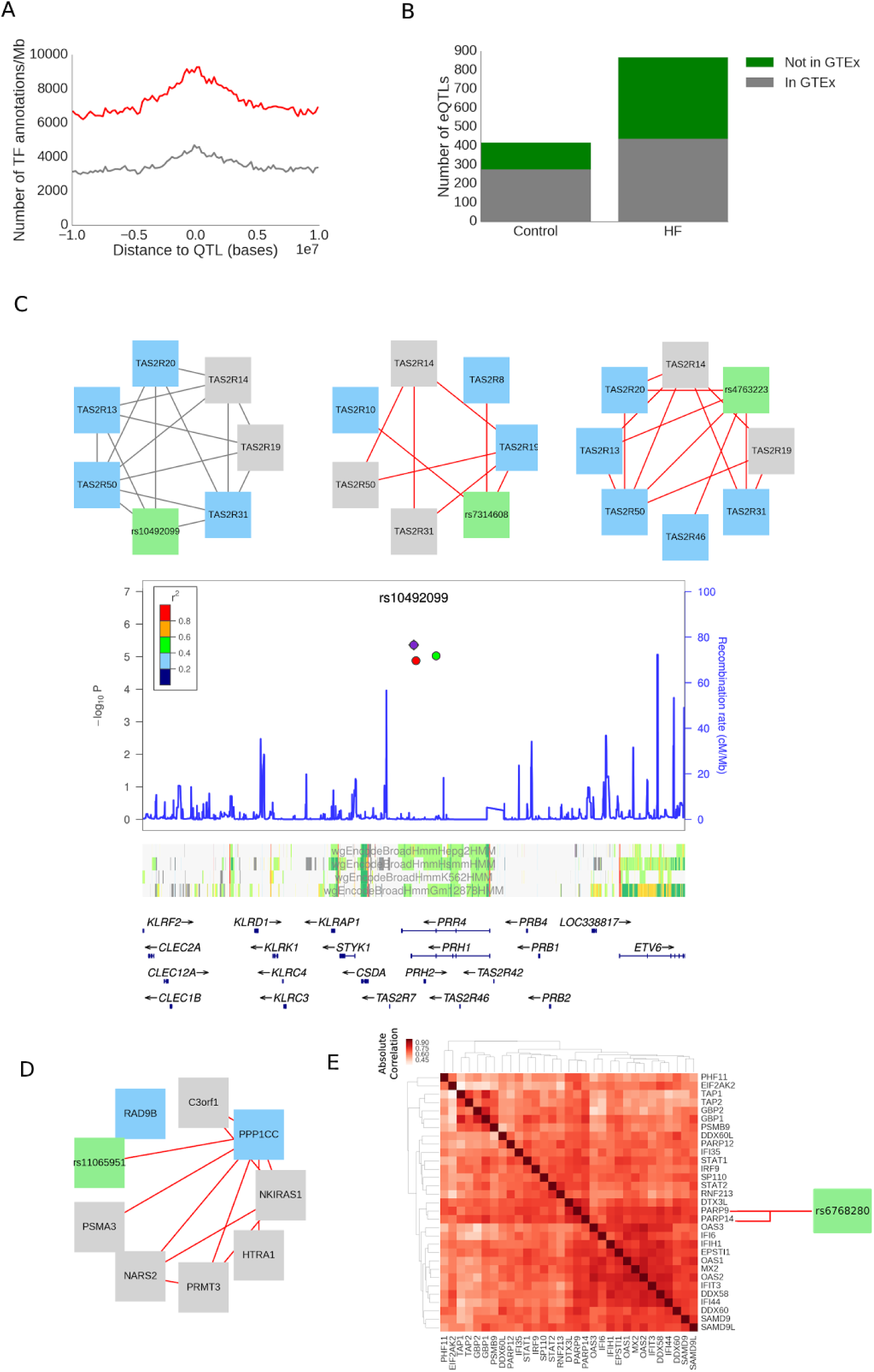
Network-enhanced discovery of heart failure eQTLs. A) Transcription factor annotation to eQTL distance distributions for failing (red) and control hearts. (B) Number of cis eQTLs found for each group that overlapped with GTEx eQTLs. (C) A set of three variants from one locus control a network of G protein coupledreceptors *TAS2R* present in both the failing and control groups. P-values indicate association significance to gene expression. (D) An eQTL that controls the proliferative function of the protein phosphate 1 through associations of *PPP1CC* and *RAD9B* (E) A variant controls immune response through association to *PARP9* and *PARP14* expression, which regulate downstream post-translational modifications of a large network of factors.

To assess the physiological impact of our eQTLs, we verified whether these loci overlapped genome-wide association study (GWAS) associations. Many of our eQTLs were replicated in several GWAS results. In particular 94 of the failing heart eQTLs replicated in the GWAS catalog ^22^ and had associations with sudden cardiac arrest, heart rate variability, and coronary heart disease among other diseases/traits (Supplementary file S3), whereas 49 of the non-failing control eQTLs had associations in the catalog, including QT interval and heart rate variability traits (Supplementary file S4).

We then sought to find modules of coordinating genetic loci and associated networks of genes within these associations by finding non-trivial connected components (i.e. with more than 3 nodes) within the bipartite association graph of variants and genes. To expand our view of the possible downstream influence of the variants in these modules, we included genes significantly connected in our networks (*r* greater than 0.7) to the genes controlled by these variants. We found 4 and 12 of these variant-gene association modules in the failing and control groups respectively (Figures S4 and S5). Notably, we found that three eQTLs, r10492099, rs7314608, and rs4763223, were within a region enriched with predicted histone modifications and controlled a network of several *TAS2R* members *in cis*, a family of G protein-coupled receptors, in both failing and control groups (Figure 3C). These associations, prevalent in both cohorts, highlight a common module of G protein-coupled receptors that have been previously observed to be expressed in the healthy and failing heart ^23,24^ and that may play a role in contraction ^25^.

Another interesting finding included rs11065951, associated with the cell cycle checkpoint gene *RAD9B* and protein phosphatase 1 catalytic subunit *PPP1CC* that were strongly connected with enzymatic genes involved in protein synthesis and proliferation in the heart failure group (Figure 3D). Protein phosphatase 1 is thought to be jointly regulating metabolic and proliferative process ^26^ and this network of associations highlights its role in heart failure ^27^. Finally, we found a large network of genes that were highly correlated with the *PARP9* and *PARP14* polymerase genes, which were associated with the rs6758280 variant in the heart failure group (Figure 3E). These polymerases have been found as regulators of the macrophage response in vascular cells ^28^. The extended association network contains several genes whose post-translational modifications are regulated by *PARP9* and *PARP14*. These include *STAT1* and *STAT2*, genes known to modulate the immune response, highlighting the up-regulation of these processes during heart failure.

In summary, our heart transplant cardiac samples and inferred gene co-expression networks enabled us to find several novel cardiac eQTLs in the failing and non-failing heart. Many of the eQTL variants were also associated with cardiac phenotypes in GWAS and some are associated with genes in highly-connected parts of the co-expression network, suggesting coordinated regulation.

### Network modeling identifies genes with differential regulatory potential in heart failure

Next, we used our networks’ topology to identify and prioritize genes that were differentially connected between the failing and control heart networks. We achieved this via two approaches: i) examining changes for each gene in its local network connectivity from control to failing networks, and ii) examining changes of a gene’s connectivity globally in known pathways. We defined local connectivity (LC) as a per gene difference in the normalized number of edges from the control to failing networks. Global connectivity (GC) was defined as the differential enrichment of known heart failure pathways in a gene’s neighborhood, comparing failing to control, with higher GC meaning a gain of pathway associations in heart failure (Supplementary Figure S6).

Using GC and LC, we assigned each gene to one of four categories. Non-hubs (N-hubs) were genes with negative LC and low GC that includes genes that lose co-expression with other genes in heart failure in comparison to control and have no change in how their neighbors are associated with known pathways. Local hubs (L-hubs) had high LC but low GC, indicating gain of gene co-expression but with genes that possessed similar pathway associations. Pathway hubs (P-hubs) were genes with high GC but negative LC, genes with a significant rewiring in pathways but losing local connectivity in the process. Finally, coordinated hubs (C-hubs) had both high GC and LC, genes that gain both connectivity and pathway function (Figure 4A).

**Figure 4.**
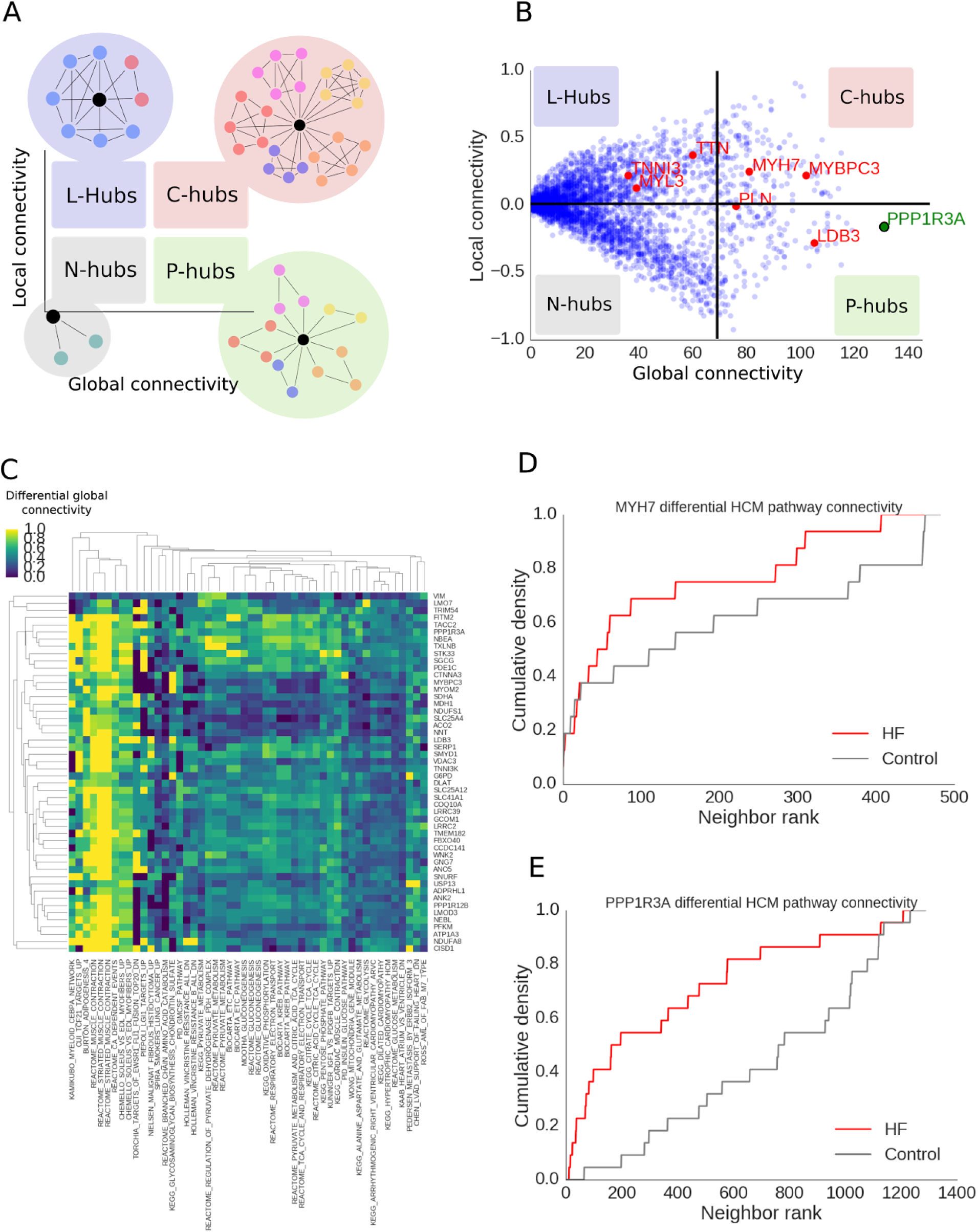
Gene prioritization through network topology. (A) Roles emerging when comparing differences in knowledge-based connectivity with local network connectivity. (B). Local and global connectivity, and roles for all genes comparing heart failure and control networks. (C) Significance statistics between top 50 differential pathways and top 50 genes with highest global connectivity (D) Difference in cumulative membership distributions for the KEGG Hypertrophic Cardiomyopathy (HCM) pathway for the myosin gene *MYH7* which is known to be involved in HCM. (E) Difference for HCM pathway enrichment in the protein phosphatase 1 regulatory subunit *PPP1R3A* is more dramatic than for *MYH7*.

We plotted gene-wise local and global connectivity metrics against each other in a scatter plot (Figure 4B) to reveal genes in these network roles (N-hubs, lower-right quadrant; L-hubs, upper-left quadrant; P-hubs, lower-right quadrants; and E-hubs, upper-right quadrant). Ranking by any of the global and local connectivity metrics gave an informative list of genes that transitioned towards centrality in the failing state and are enriched in OMIM/KEGG cardiomyopathy terms and pathways (hypertrophic and dilated cardiomyopathy KEGG pathway and OMIM terms, test p-values < 0.001). This includes the myosin heavy chain 7 *MYH7*, myosin binding protein C3 *MYBPC3*, LIM domain binding *LDB3*, and nebulette *NEBL –* genes that have previously been implicated in the Mendelian cardiac muscle diseases, hypertrophic cardiomyopathy and dilated cardiomyopathy ^29,30^ (red genes, Figure 4B). Particularly, we noted that several genes with highest differential global connectivity regulated heart failure related pathways across several processes including metabolism, muscle contraction, and cardiomyopathy-related genes (Figure 4C). In contrast, prioritization by differential gene expression did not reveal many genes genetically associated with cardiovascular disease (1 of the top 20 were associated with cardiac pathologies compared to 7 out of the top 20 for the connectivity-derived list – a significant enrichment difference [Fisher exact test p-value < 0.001]).

Among those genes whose network topology was maximally changed between failing and control human heart tissue was protein phosphatase 1 regulatory subunit 3A (*PPP1R3A*; a P-hub with high GC), which encodes a muscle-specific regulatory subunit of protein phosphatase 1 (PP1) ^31^ and has not been previously associated with heart failure. To examine the importance of *PPP1R3A* to cardiomyocyte hypertrophy across cardiomyopathic etiologies, we also examined its importance in the HCM pathway (KEGG), and found that its differential connectivity to the hypertrophic cardiomyopathy pathway (Figure 4D) exceeded even that of *MYH7* (Figure 4E), an exemplar HCM gene. Additionally, we noted that connectivity of *PPP1R3A* to our lists of sarcomeric and contraction genes was increased drastically in heart failure (Figure 1G, blue heatmap). Previous work indicates, *PPP1R3A* contains a glycogen-binding domain^32^ and is thought to promote glycogen synthesis.^33,34^ As cardiac metabolism in heart failure is known to switch toward a glucose-based metabolism, we hypothesized that *PPP1R3A* would play an important role in the transition from healthy to failing myocardium

### PPP1R3A knockdown ablates hypertrophy, a key heart failure phenotype, in vitro

To investigate the role of *PPP1R3A* in heart failure, we first determined the effect of perturbing *PPP1R3A* expression via RNA silencing on global gene expression *in vitro* (Figure 5A). We used RNA sequencing to measure global gene expression at various time points with and without *PPP1R3A* knockdown in phenylephrine treated NRVMs (an *in vitro* model of cardiomyocyte hypertrophy in heart failure). Global expression across all genes significantly changed after knockdown and phenylephrine perturbations (Student t-test < 0.05, Figure 5B). The expression of genes highly connected to *PPP1R3A* in the failing network were significantly altered and exhibited a trend towards decreasing expression (Figure 5C), evidence of a “network knockdown”. Interestingly, knockdown of *PPP1R3A* also reduced *MYH7/MYH6* gene expression ratio, a marker for cardiac hypertrophy (t-test p<0.01, Figure 5D). Cell size was also reduced (Figure 5E) as assessed by cell area (Figure 5F). Taken together, these results indicate that reduction of *PPP1R3A* expression slows cardiac hypertrophy and its associated signaling *in vitro*.

**Figure 5.**
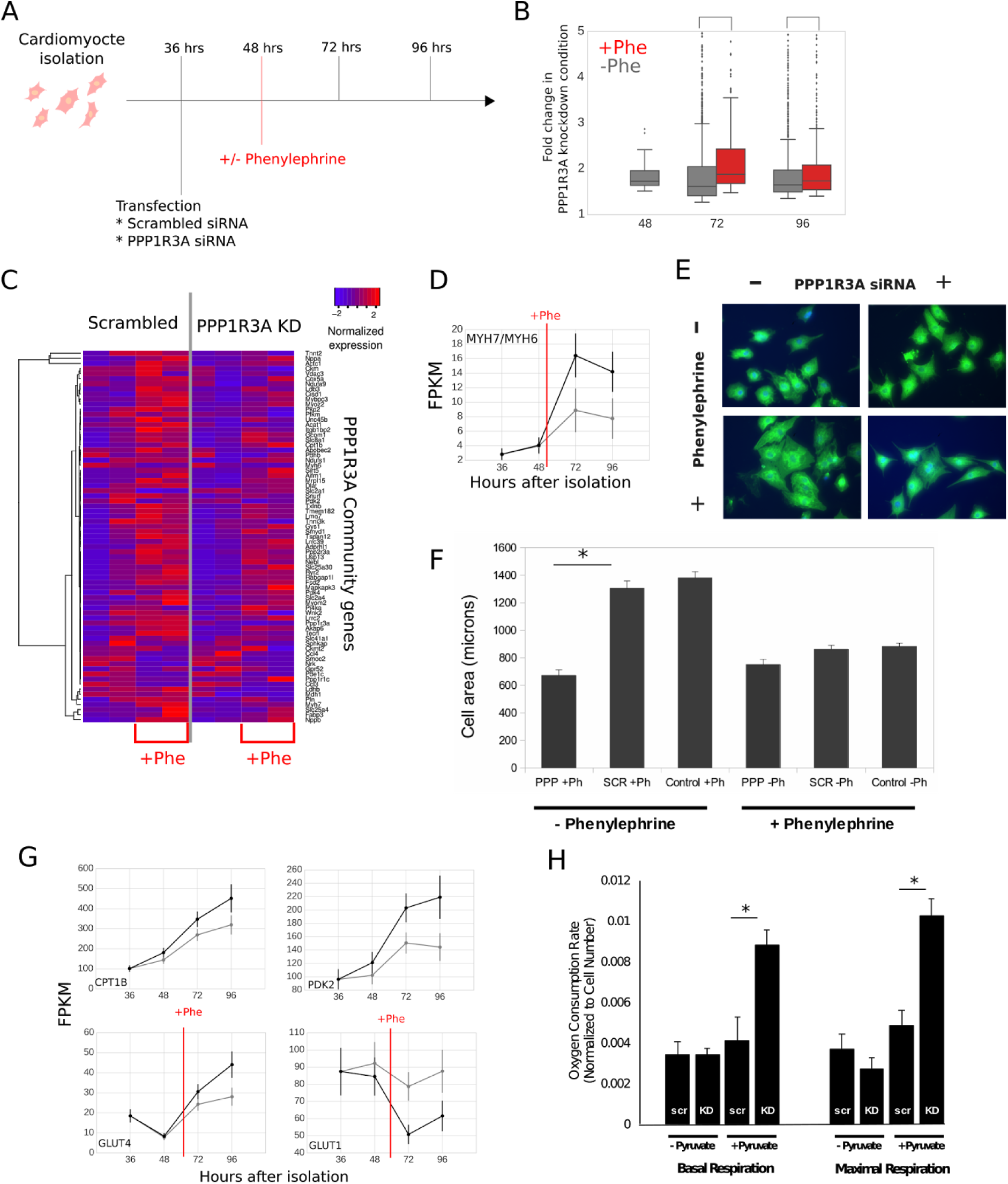
Time-course measurements of *PPP1R3A* ablation *in vitro* reveal new metabolic role. A) Experimental design and analysis pipeline for whole-transcriptome, time course measurements of an *in vitro* model of heart failure to test network-derived hypotheses. Neonatal rat ventricular myocytes (NRVMs) are isolated and split into several groups: phenylephrine treatment induces a heart failure phenotype while *PPP1R3A* siRNA transfection perturbs a highly connected gene in the human cardiac networks. As in the previous time course, RNA gene expression is measured at 36, 48, 72, and 96 hours after isolation. (B) Differences in expression fold change distributions at different timepoints after *PPP1R3A* knockdown. Clustered heatmap of normalized gene expression values (blue, lowly-expressed to red, highly-expressed) of known heart failure markers and genes in the *PPP1R3A* neighbors at several time points after isolation under hypertrophic stimulation and scrambled/*PPP1R3A* knockdown conditions. Expression level changes (X-axis) of the heart metabolic gene program as well as the *MYH7/MYH6* ratio, a marker for heart failure, as a function of time (Y-axis) and *PPP1R3A* knockdown conditions (scrambled siRNA conditions in black, *PPP1R3A* siRNA condition in gray, error bars represent standard deviations). Significant changes occur over time between *PPP1R3A* knockdown and scrambled healthy in the *PDK2* and *CPT1B* genes under normal conditions and in glucose transporters *GLUT1* and *GLUT4* under failure (phenylephrine) conditions. Further, under in vitro heart failure conditions, the marker ratio *MYH7/MYH6* is significantly decreased upon *PPP1R3A* silencing. (E) Left, cell size measurements of a sample of cells under heart failure and normal conditions, with and without *PPP1R3A* knockdown; right, sample cell images. (F) Respiratory pyruvate metabolism measured via oxygen consumption with and without *PPP1R3A* knockdown. Knockdown of PPP1R3A leads to increased basal and maximal respiratory metabolism of pyruvate as measured by oxygen consumption.

### PPP1R3A suppresses cardiomyocyte pyruvate metabolism in association with PDK expression

We hypothesized that *PPP1R3A’s* role in the development of heart failure was related to the metabolic switch from respiratory to glycolytic glucose metabolism observed in failing myocardium. We found that under phenylephrine-treated conditions, *PPP1R3A* knockdown resulted in the alteration of genes critical to glucose metabolism such as glucose transporters *GLUT1* and *GLUT4* (up-regulation and down-regulation, respectively, Figure 5G). Further, under normal cell culture conditions, knockdown of *PPP1R3A* induced a significant down-regulation of critical fatty acid metabolism genes, such as the pyruvate dehydrogenases *PDK2* and *PDK4*, and the carnitine palmitoyltransferase *CPT1B* (Figure 5G). Quantitative RT-PCR validated these findings, as treatment of NRVM with siRNA to PPP1R3A significantly decreased expression of pyruvate dehydrogenase 4 (*PDK4*) (p = 0.03, Supplemental Figure 7). *PDK4* is a major molecular driver of decreased respiratory glucose metabolism in heart failure via inactivation of pyruvate dehydrogenase ^35^. This results in shunting of pyruvate away from the oxidative TCA cycle into lactate, thus decreasing energetic efficiency of glucose metabolism in cardiomyocytes. We therefore hypothesized that decreasing *PPP1R3A* expression would lead to liberation of respiratory metabolism, and found that siRNA mediated knockdown of *PPP1R3A* leads to increased basal and maximal respiratory metabolism of pyruvate by NRVM as measured by oxygen consumption (p=0.02 (Basal Respiration) and p <0.01 (Maximal Respiration) Figure 5H, see Online Methods).

### Ppp1r3a^−/−^ mice are protected against pressure-overload-induced left ventricular failure

We then investigated the effect of *PPP1R3A* on heart failure *in vivo* using a model of pressure overload, transaortic constriction (TAC) in *Ppp1r3a*^+/+^ and *Ppp1r3a*^−/−^ mice. No difference was observed in fractional shortening between *Ppp1r3a*^+/+^ and *Ppp1r3a*^−/−^ mice at baseline. However, six and eight weeks after TAC, *Ppp1r3a*^+/+^ mice exhibited signs of heart failure with significantly reduced contractility, while *Ppp1r3a*^−/−^ mice did not (p=0.03 (6 weeks), p<0.01 (8 weeks), Figure 6A). This effect was associated with increased levels of the heart failure markers *NPPA* and *NPPB* in the LV of *Ppp1r3a*^+/+^ TAC mice, but not *Ppp1r3a*^−/−^ TAC mice (Figure 6B, compared to sham treated animals of the same genotype (*NPPA*: p=0.01 (*Ppp1r3a^+/+^*TAC vs Sham) *NPPB*: p<0.01 (*Ppp1r3a*^+/+^ TAC vs Sham). The ratio of *MYH7* to *MYH6* was increased in *Ppp1r3a^+/+^* TAC vs sham animals (p<0.01), and was not different between *Ppp1r3a*^−/−^ TAC vs Sham animals. This ratio trended toward increase in *Ppp1r3a*^−/−^ vs *Ppp1r3a*^+/+^ sham, but this difference did not reach statistical significance (p=0.08).

**Figure 6.**
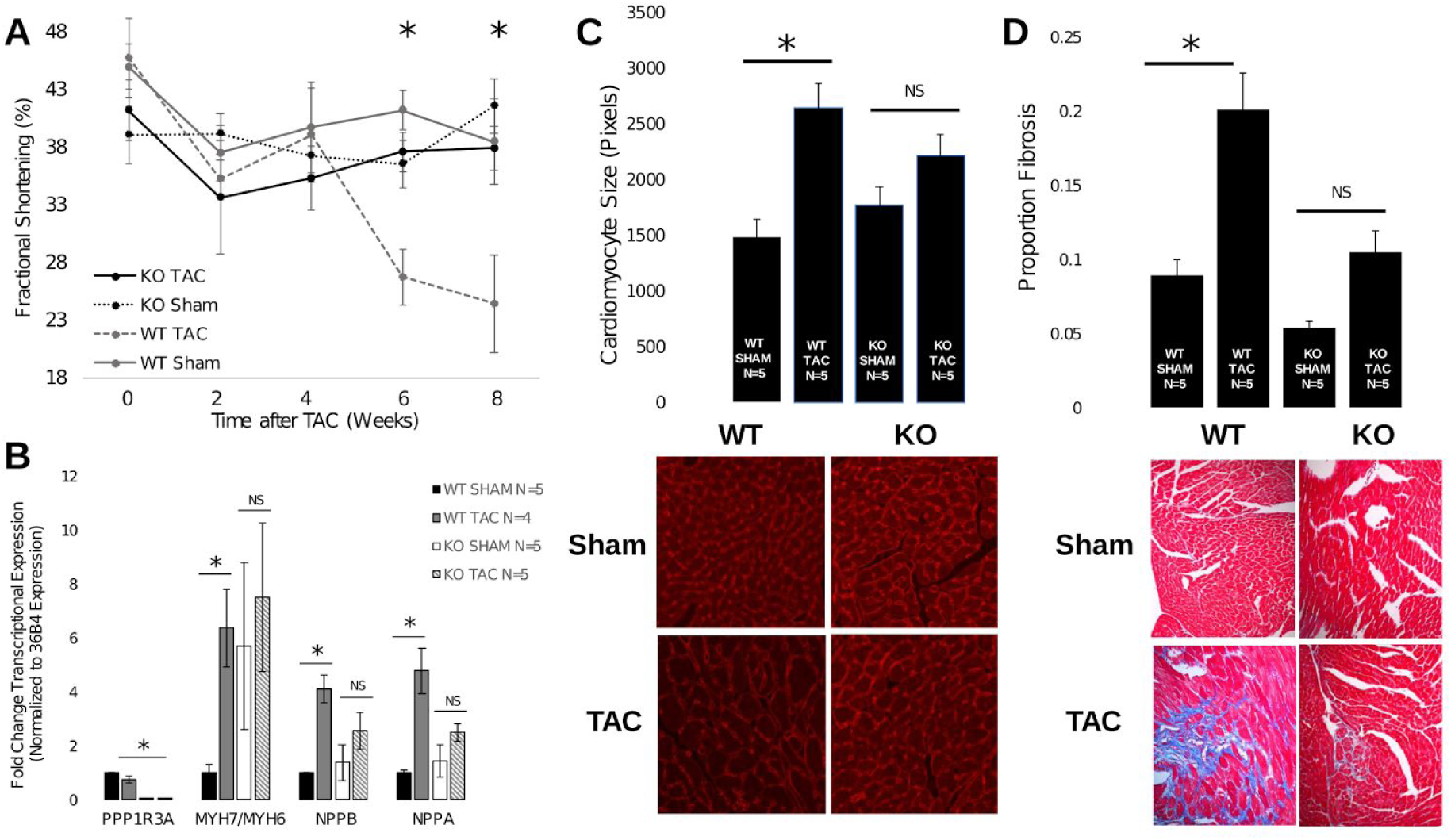
Cardiac and heart failure effects of *PPP1R3A* ablation *in vivo*. (A) Fractional shortening readings; (B) fold change gene expression of heart failure gene markers; (C) cardiomyocyte size quantification and (D) sample images; and (D) fibrosis and measurements for murine *PPP1R3A* knockdowns and wild type in an *in vivo* model of heart failure and (E) sample fibrosis images. All measurements point to an amelioration of heart failure upon *PPP1R3A* silencing.

In accordance with these findings, while immunohistochemical staining showed significantly increased cell size in *Ppp1r3a^+/+^* TAC vs sham animals (p<0.01), there was no difference between *Ppp1r3a^−/−^* TAC and sham in cardiomyocyte size, nor was there a significant difference in cardiomyocyte size between *Ppp1r3a*^+/+^ and *Ppp1r3a*^−/−^ sham-treated animals (Figure 6C). Left ventricular fibrosis as measured by trichrome staining and quantification of percent affected area showed increased fibrosis in *Ppp1r3a*^+/+^TAC mice compared to *Ppp1r3a^−/−^* TAC (p<0.01), but no significant increase in fibrosis in *Ppp1r3a*^−/−^animals after TAC compared to *Ppp1r3a*^−/−^ sham (Figure 6D).

## Discussion

We have constructed a comprehensive gene regulatory map of human heart failure. This effort has been facilitated by a systematic approach to the collection of control and failing heart tissue from the operating rooms of cardiac transplant centers and the resulting measurements have allowed us to describe several previously unrecognized molecular features of heart failure. Notably, the network structure of heart failure differs markedly from that of normal heart tissue. Specifically, we find a larger number of highly-connected regulators in the non-failing heart and a rewiring of co-expression relationships in key processes such as sarcomeric structure, excitation-contraction coupling, cell adhesion, metabolism, and cardiac remodeling -- many of these changes marking an activation and deactivation of entire subnetworks in heart failure such as Protein Phostphatase 1 related complexes and Contactin-associated proteins, respectively, indicating a complete remodeling of heart that requires coordination in a diversity of cell structures.

The inferred networks also aided in the discovery of new eQTLs in the healthy and pathological contexts. Notably, we found a greater number of eQTLs in the failing heart, half of which were novel but that were still implicated in higher phenotype associations in GWAS. In some cases, the expression of entire subnetworks of genes were found to be associated with only a few variants, such as several members of the *TAS2R* G protein coupled receptor family, receptors typically associated with the sensation of taste but recently found to be variously expressed in cardiac tissue. ^23^ Further, these eQTLs were enriched for regulatory annotations, which were more prevalent in the failing heart cohort.

Topological aspects of the networks aided in pinpointing central genes that gained and lost connectivity from non-failing to failing conditions as well as significant gain or loss of connectivity in known pathways -- measures of local and global connectivity. In particular, our network analyses identified *PPP1R3A* as a gene with a central role in heart failure. Although this gene has not previously been associated with human cardiac disease, studies in both mouse and human have found that loss-of-function mutations in *PPP1R3A* reduce glycogen content in skeletal muscle^34,36^. We found that loss of *PPP1R3A* has different effects on metabolic genes in failing versus healthy hearts *in vitro* and *in vivo*. Elimination of *PPP1R3A* in a murine model of heart failure revealed a maladaptive role for this gene in heart failure, and our *in vitro* studies implicate it in myocyte hypertrophy, and specifically in the observed metabolic switch of failing myocardium: toward inefficient glycolytic glucose metabolism and away from the use of pyruvate in respiratory metabolism.

Since genome-wide expression studies were introduced, there has been interest in quantifying genes that are significantly differentially expressed, e.g. between control and failing states. What this linear, unitary approach fails to capture are mechanisms influencing higher order phenotypes reflected in re-wiring of transcriptional partners that do not affect expression levels of specific genes. Earlier work has already led to the discovery of central genes using co-expression changes ^37,38^. Here, we expanded this use of gene co-expression by exploiting not only gene interaction degree, but also integrated topological network differences and known pathway information. In our heart failure networks, we have shown how differences between these network topology properties in failing and control can be used to highlight key genes.

## Online Methods

### Tissue collection and processing

To accelerate the understanding of the molecular underpinnings of heart failure, we established a collaborative multi-institution network with a 24/7 notification system and a team of travel-ready surgeons at major transplant centers to systematize the collection of cardiac tissue from failing hearts and unused heart transplant donors at operating rooms and remote locations. We put in place a series of best practices for procurement of explanted cardiac transplant tissue including harvesting explanted cardiac tissue at the time of cardiac surgery from subjects with heart failure undergoing transplantation and from unused donor hearts. Hearts were perfused with cold cardioplegia solution prior to cardiectomy to arrest contraction and prevent ischemic damage, and explanted cardiac tissue specimens were flash frozen in liquid nitrogen.

### Expression and genotype datasets and clinical variables

We performed RNA expression measurements and obtained genotype information in genome-wide markers for 313 patients (177 failing hearts, 136 donor, non-failing [control] hearts) using Affymetrix expression and Affymetrix Human 6.0 respectively. Clinical variables for each individual were recorded during the course of the research and were compiled using REDCap^39^.

### eQTL discovery

To test associations between gene expression in each cohort separately, we used QTLTools with an additive model accounting for gender, age, and sample site covariates. We corrected for eQTL multiple association testing using a 10000 permutations per locus in a 2 megabase window and a false discovery rate cutoff of 5%

### Quantifying global and local centrality using network and community membership parameters

The local connectivity metric (LC) of any gene *G* was calculated as the difference between max-normalized network degree of *G* considering edges exceeding an absolute correlation coefficient of 0.7. The global connectivity metric (GC) of any gene *G* was calculated as the number of gene sets that were significantly differentially enriched between gene rankings of failing and control networks obtained by ordering the genes by their absolute correlation coefficient to *G* (details included in Supplement).

### Isolation, culture, perturbation, and visualization of cardiac myocytes

Cardiac myocytes were isolated from neonatal rats using standard collagenase protocols as described previously ^9^ and cultured in serum-free, glucose-free DMEM media. In order to attenuate the effects of fibroblast contamination a final concentration of 20 μM of the fibroblast inhibitor Ara-C (Sigma-Aldrich) was incorporated. At least 1 million cells were plated in a 12-well plate, corresponding to at least 70% confluency. For phenylephrine-treated cells, 50-μM of phenylephrine was added 48 hours after isolation. For the knockdown experiments, cells were transfected either with a siRNA targeted to *PPP1R3A* (Stealth siRNA, Invitrogen) or a scrambled siRNA using the RNAiMAX system (Invitrogen) according to manufacturer's instructions; transfections were performed 24 hours after isolation. RNA extraction was performed using the Qiagen RNeasy kit according to manufacturer instructions and were DNAse-treated using the DNA-free RNA kit from Zymo research. CDNA was synthesized with the High-capacity cDNA reverse transcription kit from ABI and qRT-PCR assays were performed using KAPA SYBR FAST on a ViiA 7 ABI system.

### RNA sequencing and analysis pipeline

After RNA extraction, RNA integrity was checked using a 2100 BioAnalyzer (Agilent); all RNA samples had an RIN of 7.0 or higher. Samples were screened for *PPP1R3A* knockdown efficiency and phenylephrine treatment using qRT-PCR prior to library construction. RNA-seq libraries were prepared using the TrueSeq Stranded mRNA kit (Illumina), according to manufacturer instruction. Libraries were barcoded, quality-checked using a 2100 BioAnalyzer and run in rapid run flow cells in a HiSeq 2500 (Illumina), producing at least 30 million paired-end reads.

Sequencing reads were aligned to the *Rattus Norvegicus* rn5 UCSC reference genome using the STAR aligner ^40^. Quantification and differential expression analysis of RNA-seq data was performed using the Cufflinks package ^41^: full transcriptome assembly was performed with Cufflinks, quantified with Cuffquant, and analyzed for differential expression using Cuffdiff. All genes deemed to be significantly up or down-regulated in the main text were called as differentially expressed by Cuffdiff.

### Animals, Surgery and Phenotyping

*Ppp1r3a*^−/−^ mice (C57Bl6 background) were a generous gift from Anna de Paoli Roach ^34^. *Ppp1r3a*^+/+^animals were C57Bl6 background (Jackson). All procedures involving animal use, housing, and surgeries were approved by the Stanford Institutional Animal Care and Use Committee (IACUC). Animal care and interventions were provided in accordance with the Laboratory Animal Welfare Act.

20 male mice (10 *Ppp1r3a*^−/−^ and 10 *Ppp1r3a*^+/+^) were randomized to transaortic constriction (TAC) or sham surgery (5 in each group). Animals underwent TAC as previously described ^42^ at 10 weeks of age. Briefly, mice were anesthetized using an isoflurane inhalational chamber, intubated and ventilated. After surgical exposure of the thoracic aorta, a 6.0 silk suture was placed between the innominate and left carotid arteries to induce a constriction of ∼0.4 mm in diameter. In sham group mice, an identical procedure was conducted, without the constriction of the aorta.

*In vivo* left ventricular systolic function was evaluated by echocardiography in the short axis view as previously described ^42^. Measurements occurred at 1 day prior to surgery (baseline), 7 days and 14 days after surgery and then every 14 days prior to euthanasia and tissue collection at 8 weeks after TAC.

Upon euthanasia, heart weight, body weight and tibia length were measured by standard method (Supplementary Figure S8). Hearts were paraffin fixed, sectioned and mounted on slides. Trichrome staining as well as immunofluorescence stain for cell membrane (Rhodamine Wheat Germ Agglutinin antibody, 1:200 in PBS, Vector laboratories, Burlingame, CA) were performed for fibrosis and cell size measurements, both of which were performed using ImageJ after image capture at 20X.

### Measurement of Oxygen Consumption Rate

Freshly isolated NRVMs were plated in a 96 well plate at 75,000 cells/well and were maintained in kit medium with 0.5% fetal bovine syndrome. Transfection of siRNA to *PPP1R3A* or scrambled oligonucleotide was performed as described above 5 days after isolation. Media was changed to contain 10% fetal bovine syndrome after transfection.

Seahorse technology (XF96, Flux pack, Agilent technologies # 10-2416100) was used to measure oxygen consumption rate (OCR) 48 hours after transfection. Cells were either exposed to base media or media including pyruvate immediately before experiment. Basal metabolism was measured first. Maximal respiration was measured one minute after delivery of p-triflouromethoxyphenylhydrazone (FCCP) (uncoupler of oxidative phosphorylation). After OCR measurements were complete, viable cell number was assayed using PrestoBlue Cell Viability Assay (Thermofisher #A13261), and data were analyzed as OCR per cell.

### Data availability

Expression measurements for the human heart samples and clinical variables are available at the MAGNet portal http://www.med.upenn.edu/magnet/. Rat expression measurements are available via Amazon Web Services at http://s3.amazonaws.com/ashleylabrnaseq/timecourse_analysis

## Acknowledgements

We thank members of the Ashley lab, Stanford biomedical informatics students Winn Haynes, Albee Ling, as well as and other members of the NIH pre-print journal club for comments on the manuscript. The MAGNet heart failure study has been supported through grants for the US National Institutes of Health (HL105993, HL088577, HL098283, HL113933). This work was further supported by a predoctoral CONACyT fellowship (to PC), an NIH F32 postdoctoral fellowship (to VP) and a NIH T32 postdoctoral fellowship (to AE).

## Acknowledgements

PC, VP, AE, EA, and TC designed the study. TC and EA led data collection efforts. PC performed the computational and statistical analyses. PC, AE, and KS performed in vitro perturbation of cardiomyocytes and time course measurements. VP performed metabolic in vitro and TAC in vivo experiments. PC, AE, VP, MW, and EA wrote the manuscript with input from all authors.

